# Tumor cell-intrinsic AHR sustains IFNγ responsiveness and durable CD8 T-cell-mediated melanoma control

**DOI:** 10.64898/2026.11.19.767927

**Authors:** Susan Gellert, Metehan Celebi, Anthony C. Buzzai, Bastian Kruse, Wanyun Li, Kristin Knauth, Susanne Bonifatius, Sönke Weinert, Miriam Mengoni, Andreas D. Braun, Evelyn Gaffal, Thomas Haarmann-Stemmann, Thomas Tüting

## Abstract

The aryl hydrocarbon receptor (AHR) regulates melanoma differentiation and progression, but its tumor cell-intrinsic role in T-cell immunotherapy remains incompletely understood. Since AHR promotes inflammation-induced melanoma dedifferentiation, we hypothesized that AHR deficiency would reduce resistance to adoptive CD8 T-cell therapy (ACT) targeting melanocytic differentiation antigens. Unexpectedly, AHR-deficient HCmel12 melanomas exhibited more frequent delayed resistance to CD8 ACT despite initially impaired tumor growth. To investigate potential underlying mechanisms, we examined the responses of AHR-competent and AHR-deficient melanoma cells to prolonged interferon-γ (IFNγ) exposure. AHR-deficient cells initially exhibited enhanced IFNγ-responsive transcription but progressively lost responsiveness during chronic stimulation, accompanied by reduced MHC class I and II surface expression. Transcriptomic analyses revealed a decline in IFNγ-responsive gene expression and reduced expression of JAK/STAT signaling components, including JAK1. Treatment with the DNA methyltransferase inhibitor decitabine partially restored MHC expression, suggesting a potential contribution of epigenetic regulation. Despite their impaired responsiveness to prolonged IFNγ exposure, AHR-deficient melanoma cells remained susceptible to alternative inflammatory cytotoxic mechanisms. Importantly, CD4 ACT, which can engage broader anti-tumor effector mechanisms, eradicated AHR-deficient melanomas. Together, these findings identify an unexpected role of tumor cell-intrinsic AHR in sustaining IFNγ responsiveness during prolonged inflammatory exposure and reveal distinct outcomes of CD8- and CD4 T-cell-mediated melanoma control.

## Introduction

T-cell-directed immunotherapies have transformed the treatment of advanced melanoma (1), but therapeutic resistance remains a major challenge. Melanoma cells can evade T-cell-mediated elimination through genetic alterations and adaptive changes in their differentiation and antigen-presentation programs (2). In our previous work, we demonstrated that inflammatory signals released during adoptive CD8 T-cell therapy can induce reversible melanoma dedifferentiation, characterized by loss of melanocytic differentiation antigens and reduced recognition by antigen-specific T cells (3). We identified TNFα as a key mediator of this process, establishing inflammation-induced phenotypic plasticity as a mechanism of resistance to T-cell immunotherapy.

The aryl hydrocarbon receptor (AHR) is a ligand-activated transcription factor that integrates environmental and endogenous signals with cellular differentiation and inflammatory responses (4,5). Several studies have implicated tumor cell-intrinsic AHR signaling in melanoma progression and therapeutic resistance. In particular, Corre et al. demonstrated that sustained AHR activation promotes melanoma dedifferentiation and resistance to BRAF inhibitors, suggesting that AHR contributes to the phenotypic plasticity underlying therapeutic escape (6). However, opposing findings have also been reported, with tumor cell-intrinsic AHR restraining melanoma growth and metastatic dissemination in a different experimental model (7).

Extending these observations, we demonstrated that human melanoma cells express functional AHR and that its activation by the endogenous ligand FICZ enhances inflammation-induced dedifferentiation (8). Similar effects were observed in mouse HCmel12 melanoma cells, whereas genetic deletion of AHR reduced their metastatic dissemination following intracutaneous implantation into syngeneic mice. Together with our earlier finding that inflammation-induced dedifferentiation mediates resistance to adoptive CD8 T-cell therapy targeting melanocytic antigens (3), these observations led us to hypothesize that tumor cell-intrinsic AHR deficiency might reduce inflammation-induced dedifferentiation and thereby improve the durability of CD8 T-cell-mediated melanoma control.

Here, we investigated how tumor cell-intrinsic AHR affects the susceptibility of melanomas to adoptive T-cell immunotherapy. Contrary to our expectations, AHR deficiency compromised durable CD8 T-cell-mediated tumor control. We found that AHR-deficient melanoma cells initially mounted an enhanced transcriptional response to IFNγ but progressively lost their responsiveness during prolonged exposure, accompanied by reduced expression of JAK/STAT signaling components and antigen-presentation machinery. Nevertheless, these melanomas remained susceptible to alternative cytotoxic mechanisms including CD4 T-cell immunotherapy. Our findings identify a previously unrecognized role of tumor cell-intrinsic AHR in maintaining IFNγ responsiveness during prolonged inflammatory exposure and reveal differences in the susceptibility of AHR-deficient melanomas to CD8- and CD4 T-cell-mediated tumor control.

## Results

### Tumor cell-intrinsic AHR deficiency compromises durable CD8 T-cell-mediated melanoma control

To investigate whether tumor cell-intrinsic expression of AHR affects the response to T-cell immunotherapy, we used HCmel12 AHR-KO mouse melanoma cells previously generated by CRISPR/Cas9-mediated gene targeting (8). We first evaluated their response to an adoptive CD8 T-cell therapy protocol (CD8 ACT; Fig. 1a) developed in our laboratory (3,9). Both AHR-expressing HCmel12 CRISPR-control and AHR-KO melanoma cells grew progressively with similar kinetics upon intracutaneous implantation into syngeneic C57BL/6 mice. Treatment of tumors reaching a diameter of 3–4 mm with CD8 ACT initially impaired progressive growth in all mice. Eventually, 5 of 12 mice inoculated with AHR-competent melanoma cells remained tumor-free at the end of the observation period. Contrary to our expectations, only 2 of 20 mice bearing AHR-deficient melanomas achieved durable tumor control (Fig. 1b,c). Notably, the tumor growth dynamics revealed a higher frequency of delayed resistance to CD8 ACT in AHR-deficient compared with AHR-competent melanomas, suggesting that tumor cell-intrinsic AHR contributes to durable CD8 T-cell-mediated melanoma control.

**Figure 1.**
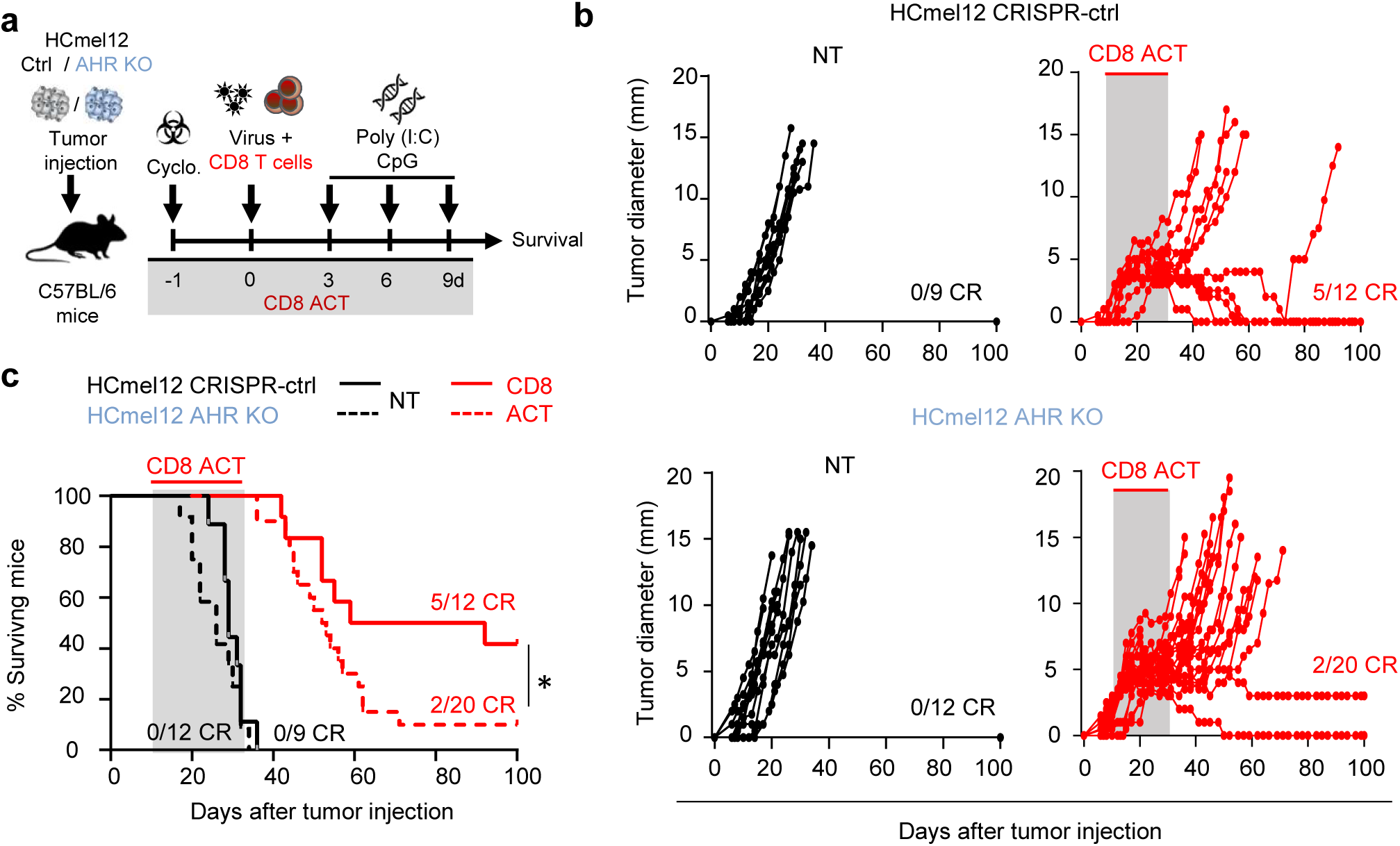
Tumor-intrinsic AHR loss promotes delayed escape from CD8 ACT. **a**, Experimental protocol for CD8 adoptive T-cell therapy (CD8 ACT) of established HCmel12 CRISPR-control and AHR-knockout (AHR-KO) melanomas in C57BL/6 mice. Treatment was initiated when tumors reached a diameter of approximately 3-4 mm, typically 10-14 days after tumor inoculation. Mice received cyclophosphamide preconditioning one day before adoptive transfer of Pmel-1 TCR-transgenic CD8+ T cells and adenoviral vaccination with AdPT, followed by intratumoral injections of the innate immune stimuli poly(I:C) and CpG. **b**, Individual tumor growth curves of mice bearing HCmel12 CRISPR-control or HCmel12 AHR-KO melanomas, either left untreated (NT) or treated with CD8 ACT. Tumor growth is shown relative to tumor inoculation (day 0). The gray shading indicates the approximate CD8 ACT treatment period. CR, Complete responders. **c**, Kaplan-Meier survival curves of mice bearing HCmel12 CRISPR-control or AHR-KO melanomas, either left untreated or treated with CD8 ACT. Survival is shown relative to tumor inoculation and was compared using the log-rank (Mantel–Cox) test. \**P* ≤ 0.05. CR, Complete responders.

### AHR-deficient melanoma cells progressively lose MHC expression during chronic IFNγ exposure

Interferon gamma (IFNγ) is a major effector cytokine secreted by activated CD8 T cells, and tumor cell-intrinsic IFNγ signaling is important for T-cell-mediated tumor control. Given previous evidence that AHR interacts with STAT1 and modulates IFN signaling (10,11), and that prolonged IFNγ exposure can alter melanoma susceptibility to immunotherapy (12), we investigated whether AHR deficiency affects melanoma cell responsiveness to chronic IFNγ stimulation. To mimic an important component of persistent immune pressure during immunotherapy *in vivo*, we treated HCmel12 CRISPR-control and AHR-KO melanoma cells with IFNγ twice weekly for up to 28 days and monitored MHC class I (MHC-I) and class II (MHC-II) surface expression longitudinally by flow cytometry (Fig. 2a). Both cell populations initially upregulated MHC-I expression, with comparable responses after three days. By day 7, MHC-I expression began to decline in AHR-KO cells, followed by a pronounced reduction at day 14 that persisted until day 28. In contrast, CRISPR-control cells maintained robust MHC-I expression throughout the experiment (Fig. 2b,c top panels). MHC-II expression followed a similar pattern, with comparable initial induction in both cell populations, followed by progressive loss in HCmel12 AHR-KO cells but sustained expression in CRISPR-control cells (Fig. 2b,c bottom panels). These findings indicate that AHR deficiency selectively compromises the maintenance rather than the initial induction of IFNγ-dependent MHC expression in melanoma cells.

**Figure 2.**
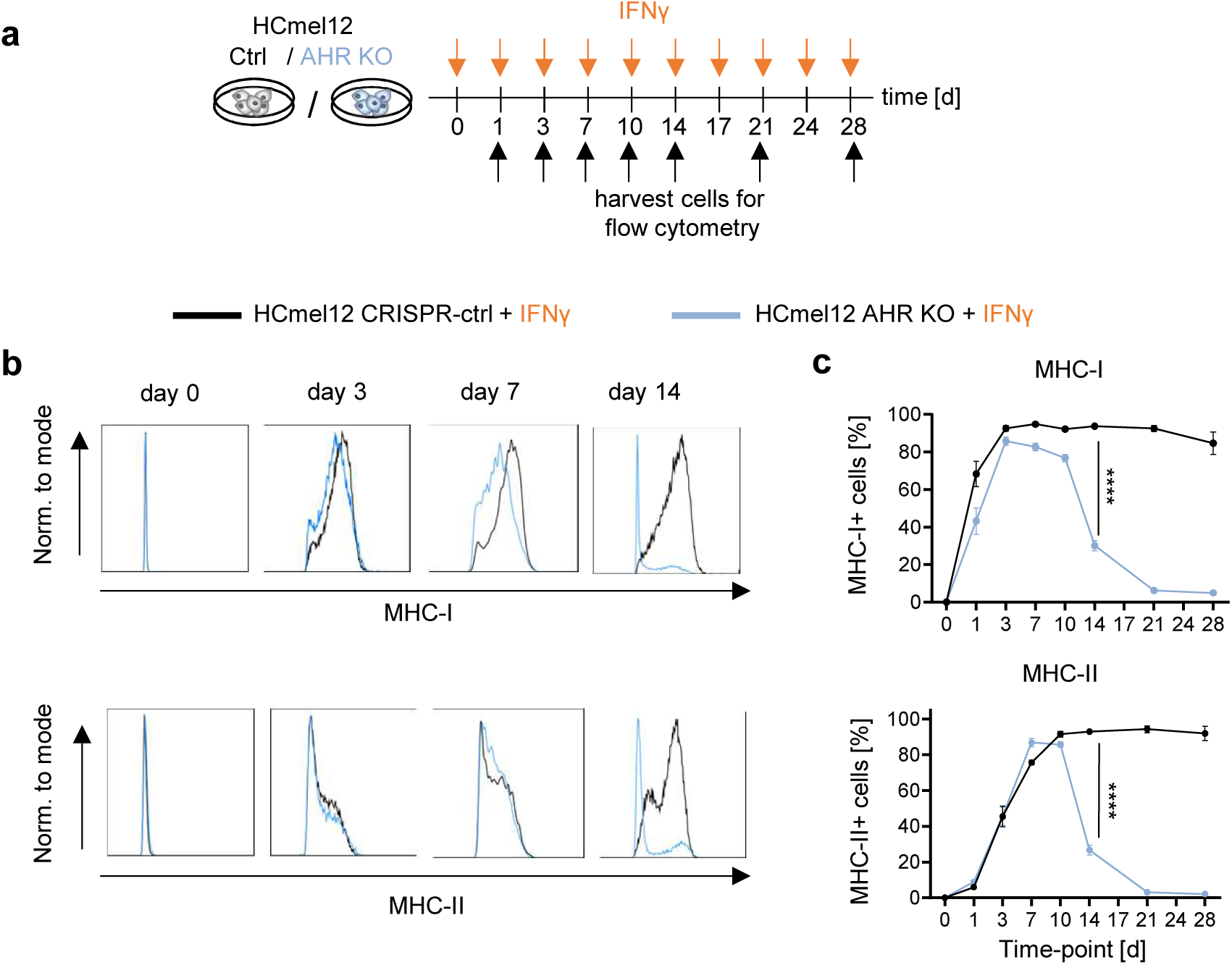
AHR-deficient melanoma cells progressively lose MHC expression during chronic IFNγ exposure. **a**, Experimental protocol for chronic IFNγ stimulation of HCmel12 CRISPR-control and AHR-KO melanoma cells. Cells were repeatedly stimulated with IFNγ (1,000 U/ml) for up to 28 days and harvested at the indicated time points for flow cytometric analysis. **b**, Representative flow cytometric histograms showing MHC-I (top) and MHC-II (bottom) surface expression on HCmel12 CRISPR-control (black) and AHR-KO (blue) melanoma cells before (day 0) and after 3, 7, and 14 days of IFNγ exposure. **c**, Percentage of MHC-I-positive (top) and MHC-II-positive (bottom) cells during chronic IFNγ exposure. Data are presented as mean ± SD from 2 independent experiments. Statistical significance was determined by Two-Way ANOVA followed by Tukey’s multiple-comparisons test. \*\*\*\**P* ≤ 0.0001.

### AHR deficiency alters the temporal dynamics of IFNγ-induced transcriptional programs in melanoma cells

To investigate whether the progressive loss of IFNγ-induced MHC expression in AHR-deficient melanoma cells was associated with broader transcriptional changes, we performed longitudinal bulk RNA sequencing of HCmel12 CRISPR-control and AHR-KO cells after 0, 3, 7, and 14 days of repeated IFNγ stimulation (Fig. 3a). We identified 1,741 differentially expressed genes across the experimental conditions and used unsupervised k-means clustering to characterize their temporal expression patterns (Fig. 3b). This analysis revealed seven distinct clusters (C1–C7) with different expression dynamics in AHR-competent and AHR-deficient melanoma cells (Fig. 3b,c). Notably, genes in clusters C1 and C2 exhibited sustained expression in HCmel12 CRISPR-control cells, whereas their expression in AHR-KO cells was initially enhanced but declined during prolonged IFNγ exposure. To identify the biological processes associated with these transcriptional patterns, we performed enrichment analysis using MSigDB Hallmark gene sets. This revealed enrichment of inflammatory response-associated programs, including interferon signaling, particularly in clusters C1 and C2 (Fig. 3d). Other clusters showed distinct temporal patterns, indicating that AHR deficiency did not uniformly suppress transcriptional responses during chronic IFNγ stimulation. These findings suggest that tumor cell-intrinsic AHR contributes to the maintenance of distinct inflammatory transcriptional programs during prolonged IFNγ exposure.

**Figure 3.**
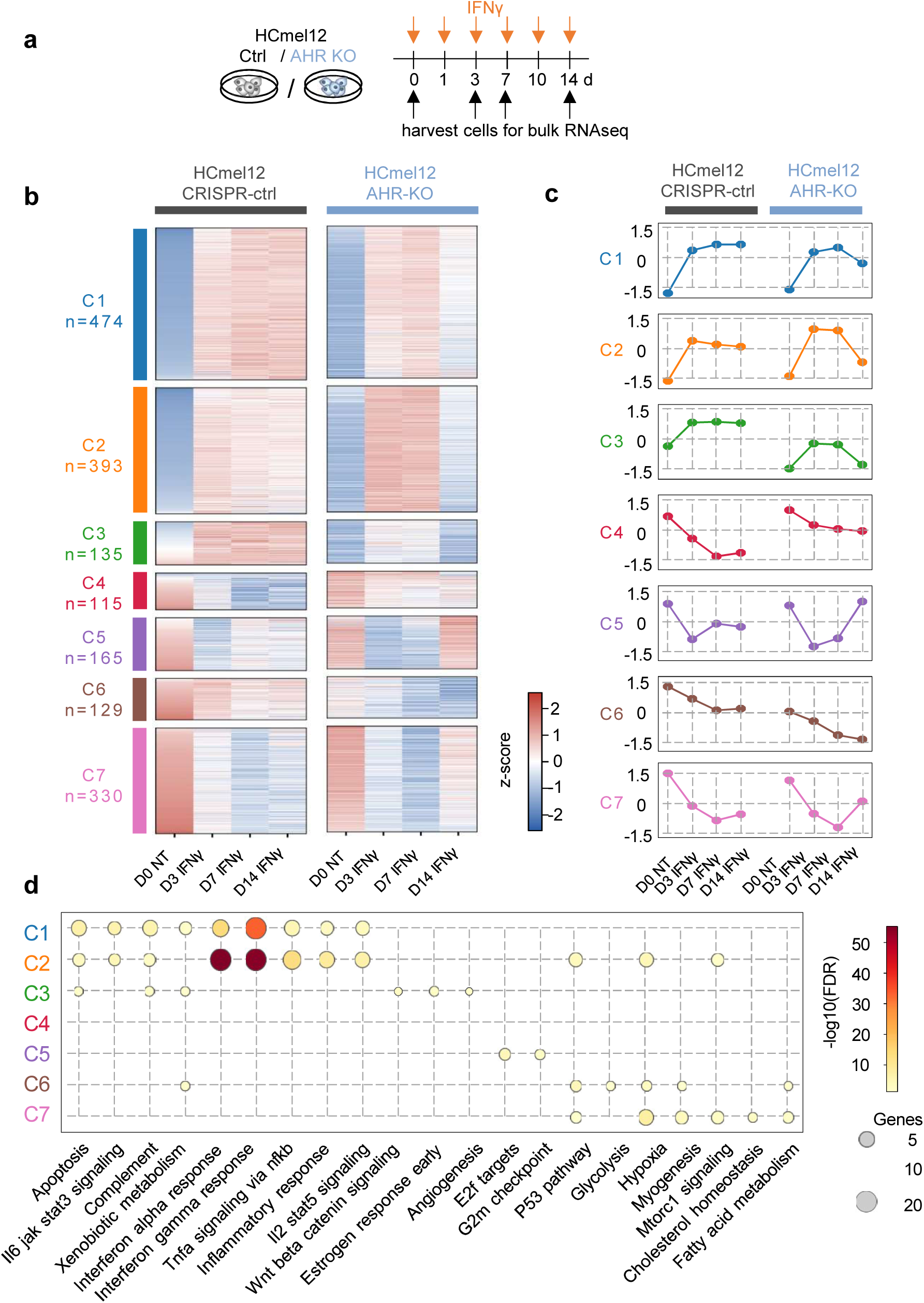
AHR deficiency alters the temporal dynamics of IFNγ-induced transcriptional programs in melanoma cells. **a**, Experimental protocol for chronic IFNγ stimulation of HCmel12 CRISPR-control and AHR-KO melanoma cells. Cells were repeatedly exposed to IFNγ and harvested at days 0, 3, 7, and 14 for bulk RNA sequencing. **b**, Heatmaps showing the expression patterns of 1,741 differentially expressed genes (DEGs) in HCmel12 CRISPR-control and AHR-KO melanoma cells before (day 0) and after 3, 7, and 14 days of IFNγ exposure. Genes were selected based on differential expression (adjusted *P* < 0.05, |log₂FC| ≥ 1) and a mean normalized count ≥ 50 in at least one experimental condition. Unsupervised k-means clustering identified seven distinct temporal expression patterns (C1–C7). Colors represent gene-wise Z-scores calculated from mean log₂-transformed DESeq2-normalized counts across all eight experimental conditions (n = 3 biological replicates per condition). **c**, Mean Z-score profiles (cluster centroids) showing the temporal expression dynamics of each cluster in HCmel12 CRISPR-control (left) and AHR-KO (right) cells. Colors correspond to the clusters shown in **b**. **d**, Over-representation analysis of MSigDB Hallmark gene sets within each cluster using a one-sided hypergeometric test with all protein-coding genes tested in the differential expression analysis as the background and Benjamini–Hochberg correction for multiple testing. The union of the six most significantly enriched gene sets per cluster is shown. Dots indicate significant enrichment (FDR < 0.05), with dot size representing the number of overlapping genes and color indicating −log₁₀(FDR).

### AHR-deficient melanoma cells fail to sustain IFNγ-responsive transcription and antigen-presentation programs

To characterize the inflammatory transcriptional programs affected by AHR deficiency in greater detail, we first analyzed the murine MSigDB Hallmark IFNγ-response gene set. At baseline, HCmel12 CRISPR-control and AHR-KO cells exhibited largely similar expression patterns. Following IFNγ stimulation, both cell populations induced IFNγ-responsive genes, but AHR-KO cells displayed a stronger transcriptional response at days 3 and 7. Strikingly, this enhanced response was not sustained. By day 14, expression of the IFNγ-response gene set had declined markedly in HCmel12 AHR-KO cells, whereas CRISPR-control cells maintained robust expression throughout the treatment period (Fig. 4a–c). Given the importance of IFNγ for tumor antigen presentation, we next examined the expression of genes involved in MHC-I antigen processing and presentation. Similar to the broader IFNγ-response signature, these genes were initially induced in both cell populations but showed a pronounced decline in HCmel12 AHR-KO cells during prolonged IFNγ exposure, while their expression remained sustained in CRISPR-control cells (Fig. 4d,e). A comparable pattern was observed for MHC-II-associated genes, including *Ciita* and MHC-II components, which also exhibited reduced expression in AHR-KO cells at day 14 (Fig. 4f,g). Together, these findings indicate that AHR deficiency enhances the early transcriptional response to IFNγ but compromises its long-term maintenance, including the expression of genes involved in MHC-I and MHC-II antigen presentation.

**Figure 4.**
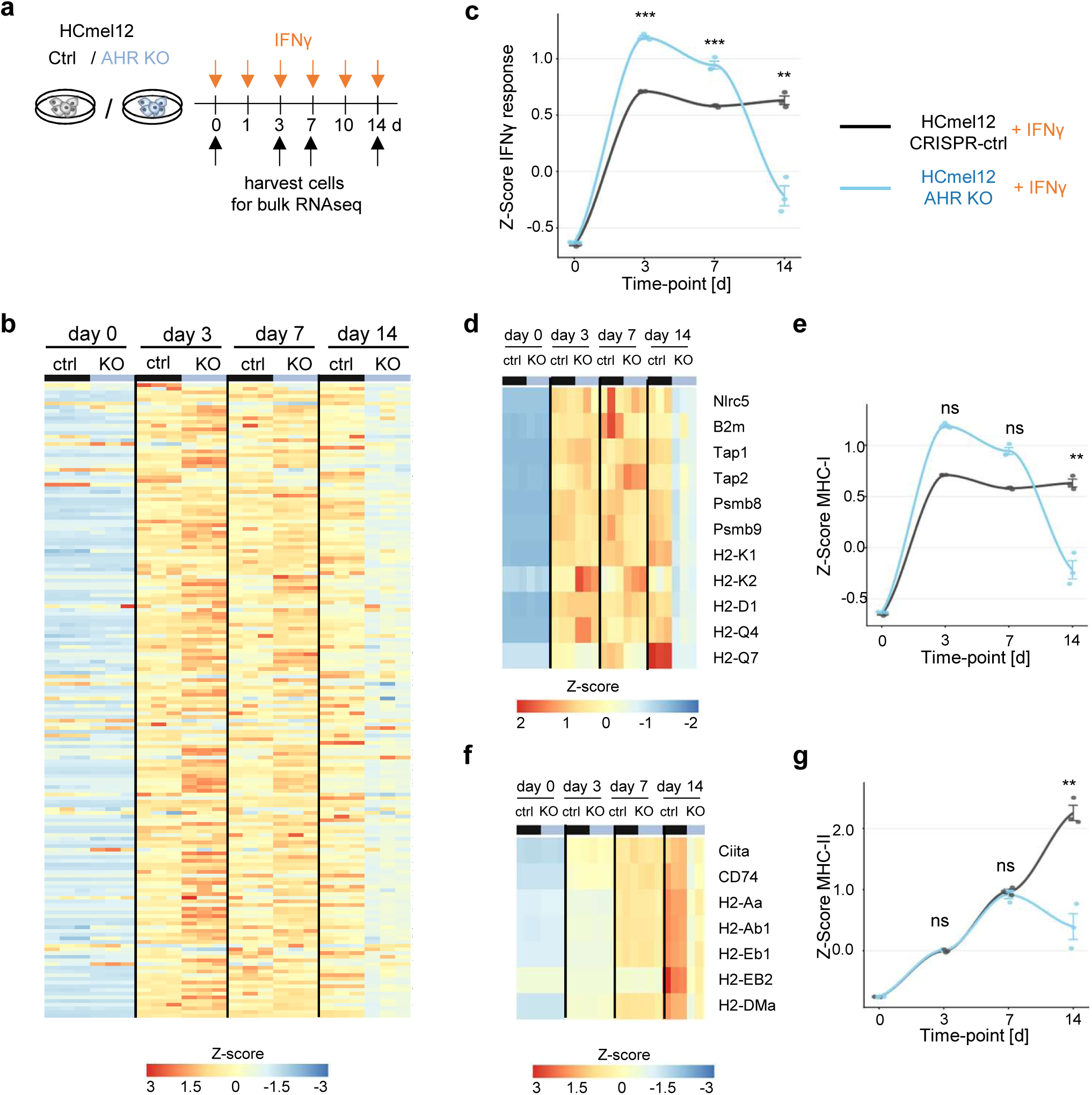
AHR-deficient melanoma cells fail to sustain IFNγ-responsive transcription and antigen-presentation programs. **a**, Experimental protocol for chronic IFNγ stimulation of HCmel12 CRISPR-control and AHR-KO melanoma cells. Cells were repeatedly exposed to IFNγ and harvested at days 0, 3, 7, and 14 for bulk RNA sequencing. **b**, Heatmap showing the expression of genes belonging to the murine MSigDB Hallmark IFNγ-response gene set in HCmel12 CRISPR-control and AHR-KO melanoma cells during chronic IFNγ exposure. Colors indicate standardized gene expression (Z-scores). **c**, Temporal changes in the mean expression Z-score of the Hallmark IFNγ-response gene set in HCmel12 CRISPR-control (black) and AHR-KO (blue) cells. **d, e**, Heatmap (**d**) and corresponding temporal expression profiles (**e**) of genes involved in MHC-I antigen processing and presentation. **f, g**, Heatmap (**f**) and corresponding temporal expression profiles (**g**) of genes involved in MHC-II antigen processing and presentation. Data are presented as mean ± SD from three biological replicates per condition. Statistical significance between HCmel12 CRISPR-control and AHR-KO cells at individual time points was determined using Welch’s two-sample *t*-test. ns, not significant; **P ≤ 0.01; ***P ≤ 0.001.

### Loss of sustained IFNγ responsiveness in AHR-deficient melanoma cells is associated with downregulation of JAK/STAT signaling components

To investigate whether the progressive decline in IFNγ-responsive transcription was associated with changes in the signaling machinery itself, we examined the expression of key components of the IFNγ receptor–JAK/STAT pathway during chronic IFNγ exposure (Fig. 5a,b). Analysis of a selected set of pathway-associated genes revealed initially increased expression in both HCmel12 CRISPR-control and AHR-KO cells. HCmel12 AHR-KO cells then showed a pronounced decline after prolonged IFNγ stimulation. In contrast, HCmel12 CRISPR-control cells maintained relatively stable expression throughout the treatment period (Fig. 5c). Analysis of individual pathway components revealed distinct expression patterns. The IFNγ receptor genes *Ifngr1* and *Ifngr2* remained relatively stable in both cell populations, whereas *Jak1*, *Jak2*, *Stat1*, *Irf1*, and *Socs1* showed progressively reduced expression in HCmel12 AHR-KO cells, particularly at day 14. Among these genes, *Jak1* exhibited a particularly pronounced decline, with expression falling below baseline levels before IFNγ stimulation (Fig. 5d). Consistent with these transcriptional changes, Western blot analysis revealed reduced JAK1 protein expression in HCmel12 AHR-KO cells after 14 days of IFNγ exposure compared with CRISPR-control cells (Fig. 5e).

**Figure 5.**
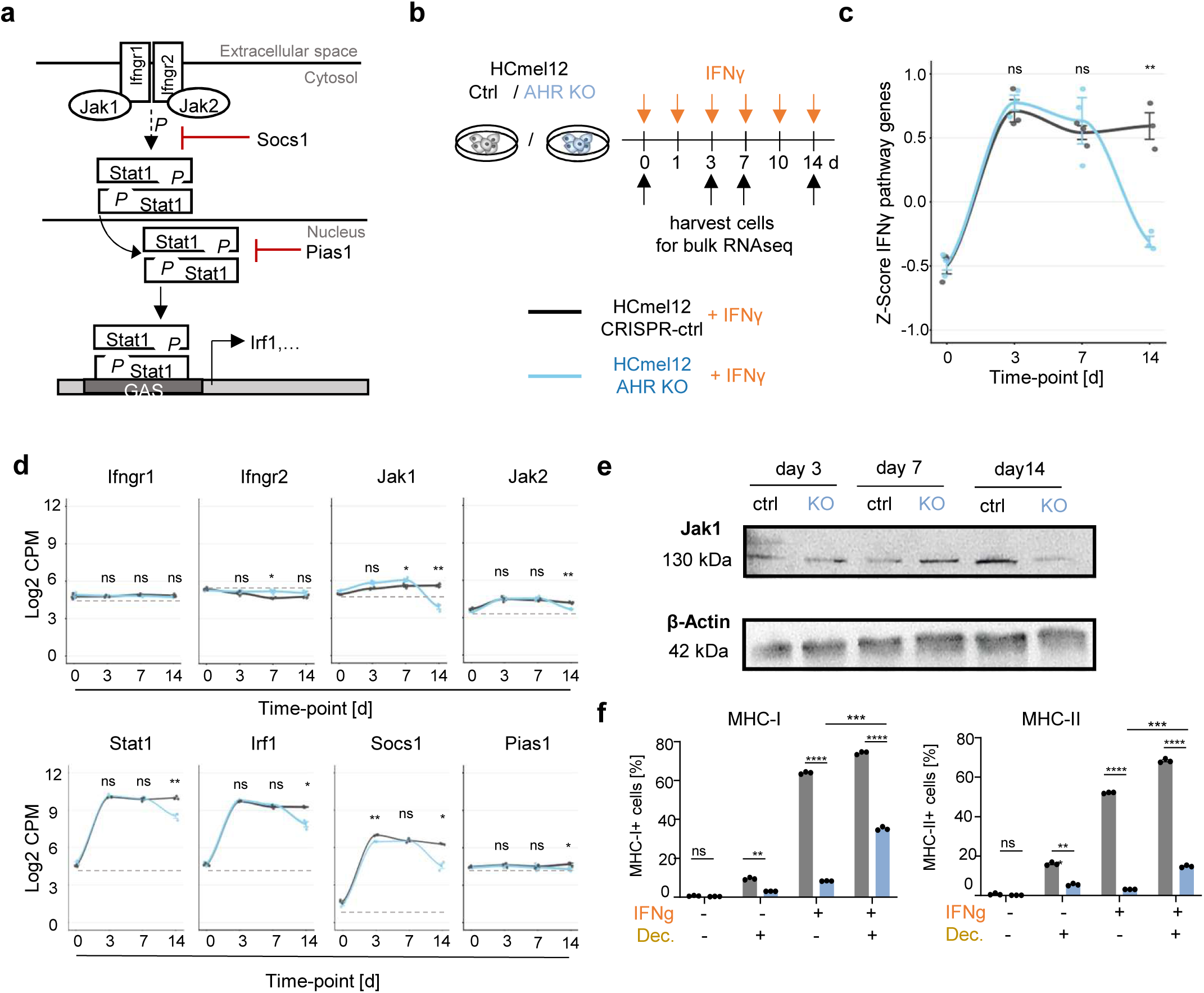
Chronic IFNγ exposure is associated with downregulation of JAK/STAT signaling components in AHR-deficient melanoma cells. **a**, Schematic representation of the canonical IFNγ receptor-JAK/STAT signaling pathway, including the negative regulators SOCS1 and PIAS1. **b**, Experimental protocol for chronic IFNγ stimulation of HCmel12 CRISPR-control and AHR-KO melanoma cells. Cells were repeatedly exposed to IFNγ and harvested at days 0, 3, 7, and 14 for bulk RNA sequencing. **c**, Temporal changes in the mean expression Z-score of selected IFNγ signaling pathway genes (*Ifngr1, Ifngr2, Jak1, Jak2, Stat1, Irf1, Socs1,* and *Pias1*) in HCmel12 CRISPR-control (black) and AHR-KO (blue) cells. **d**, Temporal expression profiles of individual IFNγ signaling pathway genes in HCmel12 CRISPR-control and AHR-KO cells, shown as log₂-transformed Count Per Million (CPM). Data are presented as mean ± SD from three biological replicates per condition. Statistical significance between the two cell populations at individual time points was determined using Welch’s two-sample *t*-test. **e**, Representative Western blot showing JAK1 protein expression in HCmel12 CRISPR-control and AHR-KO cells after 3, 7, and 14 days of IFNγ exposure. β-Actin served as a loading control. **f**, Percentage of MHC-I-positive (left) and MHC-II-positive (right) melanoma cells 4 days after application of decitabine on day 28 of chronic IFNγ exposure. Data are presented as mean ± SD. Statistical significance was determined by Two-Way ANOVA followed by Tukey’s multiple-comparisons test. ns, not significant; \**P* ≤ 0.05; \*\**P* ≤ 0.01; *\*\*\*P* ≤ 0.001, *\*\*\*\*P* ≤ 0.0001.

We hypothesized that the progressive loss of IFNγ responsiveness in AHR-deficient melanoma cells might involve epigenetic changes that accumulate during prolonged inflammatory stimulation. To explore a potential contribution of DNA methylation, we treated HCmel12 CRISPR-control and AHR-KO cells with the DNA methyltransferase inhibitor decitabine following 28 days of repeated IFNγ exposure. Decitabine partially restored MHC-I and MHC-II expression in HCmel12 AHR-KO cells, whereas MHC expression remained largely unchanged in CRISPR-control cells (Fig. 5f). Together, these findings indicate that the progressive loss of IFNγ responsiveness in AHR-deficient melanoma cells is associated with downregulation of intracellular JAK/STAT signaling components, particularly JAK1, and that the acquired reduction in MHC expression is partially reversible.

### AHR-deficient melanomas remain susceptible to alternative cytotoxic mechanisms and CD4 T-cell immunotherapy

In our previous work, we demonstrated that CD4 T-cell immunotherapy can eradicate IFNγ-unresponsive HCmel12 *Jak1*-KO melanomas that resist CD8 ACT through an indirect cytotoxic mechanism involving nitric oxide-producing myeloid cells (13). Given the progressive loss of IFNγ responsiveness in HCmel12 AHR-KO melanoma cells, we hypothesized that these tumors might remain susceptible to CD4 ACT despite their reduced responsiveness to CD8 T-cell immunotherapy. We first investigated whether chronic IFNγ exposure altered the susceptibility of HCmel12 CRISPR-control and AHR-KO cells to inflammatory cytotoxicity. Cells were either left untreated or exposed to IFNγ for 28 days, followed by stimulation with IFNγ and TNFα or the nitric oxide donor SNAP in combination with TNFα for 24 hours. Cell death was assessed by Annexin V/propidium iodide staining (Fig. 6a). Without prior IFNγ exposure, both cell populations were susceptible to IFNγ/TNFα- and SNAP/TNFα-mediated cytotoxicity (Fig. 6b,d). Following chronic IFNγ exposure, however, HCmel12 AHR-KO cells exhibited markedly reduced susceptibility to IFNγ/TNFα-induced cell death, whereas CRISPR-control cells remained sensitive (Fig. 6c). In contrast, SNAP/TNFα induced substantial cell death in both cell populations even after prolonged IFNγ exposure (Fig. 6e). We next investigated the efficacy of CD4 ACT against established HCmel12 CRISPR-control and AHR-KO melanomas in syngeneic C57BL/6 mice (Fig. 6f). Whereas 4 of 8 mice inoculated with AHR-competent melanoma cells achieved durable tumor control, 8 of 9 mice inoculated with AHR-deficient melanoma cells remained tumor-free at the end of the observation period (Fig. 6g,h). Together, these findings demonstrate that AHR-deficient melanoma cells retain susceptibility to nitric oxide-associated cytotoxicity despite their reduced responsiveness to IFNγ/TNFα and that CD4 T-cell immunotherapy can achieve durable control of AHR-deficient melanomas *in vivo*.

**Figure 6.**
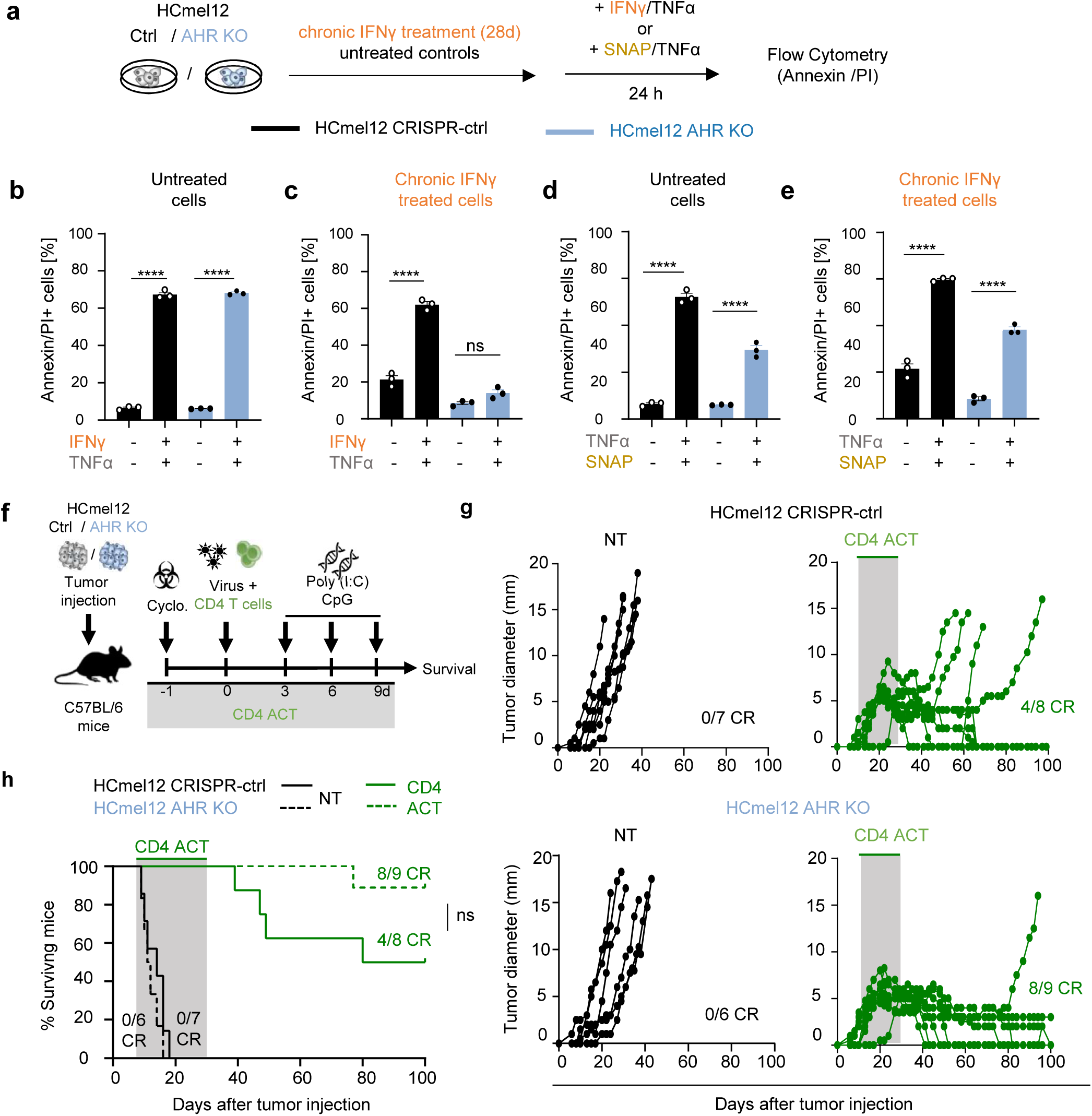
AHR-deficient melanoma cells retain susceptibility to nitric oxide-mediated cytotoxicity and CD4 T-cell immunotherapy. **a**, Experimental protocol for cell death assays in HCmel12 CRISPR-control and AHR-KO melanoma cells. Cells were either left untreated or repeatedly stimulated with IFNγ for 28 days, followed by treatment with IFNγ/TNFα or the nitric oxide donor SNAP in combination with TNFα for 24 hours. Cell death was assessed by Annexin V/propidium iodide (PI) staining and flow cytometry. **b, c**, Percentage of Annexin V/PI-positive HCmel12 CRISPR-control (**b**) and AHR-KO (**c**) melanoma cells following IFNγ/TNFα treatment, either without prior IFNγ exposure or after 28 days of chronic IFNγ stimulation. **d, e**, Percentage of Annexin V/PI-positive HCmel12 CRISPR-control (**d**) and AHR-KO (**e**) melanoma cells following SNAP/TNFα treatment, either without prior IFNγ exposure or after 28 days of chronic IFNγ stimulation. **f**, Experimental protocol for CD4 adoptive T-cell therapy (CD4 ACT) of established HCmel12 CRISPR-control and AHR-KO melanomas in C57BL/6 mice. Mice received cyclophosphamide preconditioning followed by adoptive transfer of tumor-reactive CD4+ T cells, adenoviral vaccination, and intratumoral administration of poly(I:C) and CpG. **g**, Individual tumor growth curves of mice bearing HCmel12 CRISPR-control or AHR-KO melanomas, either left untreated (NT) or treated with CD4 ACT. Tumor growth is shown relative to tumor inoculation (day 0). The gray shading indicates the approximate CD4 ACT treatment period. CR, Complete responders. **h**, Kaplan–Meier survival curves of mice bearing HCmel12 CRISPR-control or AHR-KO melanomas, either left untreated or treated with CD4 ACT. Survival is shown relative to tumor inoculation and was compared using the log-rank (Mantel–Cox) test. Statistical analyses of cell death were performed using One-Way ANOVA followed by Tukey‘s multiple comparisons test. ns, not significant; \*\*\*\**P* ≤ 0.0001. CR, Complete responders.

### CD4 ACT promotes infiltration of iNOS-expressing myeloid populations into melanomas

To investigate whether CD4 ACT induces the recruitment of myeloid effector cells in AHR-deficient melanomas, we analyzed the immune infiltrates of HCmel12 CRISPR-control and AHR-KO tumors seven days after CD4 T-cell transfer using spectral flow cytometry (Fig. 7a). CD4 ACT markedly increased the numbers of tumor-infiltrating immune cells in both groups, particularly monocytes/macrophages and neutrophils. Notably, the accumulation of myeloid cells appeared more pronounced in AHR-KO than in CRISPR-control melanomas (Fig. 7b). We next examined the expression of inducible nitric oxide synthase (iNOS) in these myeloid populations. CD4 ACT increased the proportions of iNOS-expressing neutrophils and Ly6C^high^ monocytes/macrophages in both tumor types. Interestingly, AHR-KO melanomas exhibited a higher proportion of iNOS-expressing Ly6C^high^ monocytes/macrophages following CD4 ACT, whereas the proportion of iNOS-expressing neutrophils was lower than in CRISPR-control melanomas (Fig. 7c). Together, these findings indicate that CD4 ACT promotes the infiltration of iNOS-expressing myeloid cells into AHR-deficient melanomas, providing a potential cellular basis for their susceptibility to CD4 T-cell-mediated tumor eradication despite impaired sustained IFNγ responsiveness.

**Figure 7.**
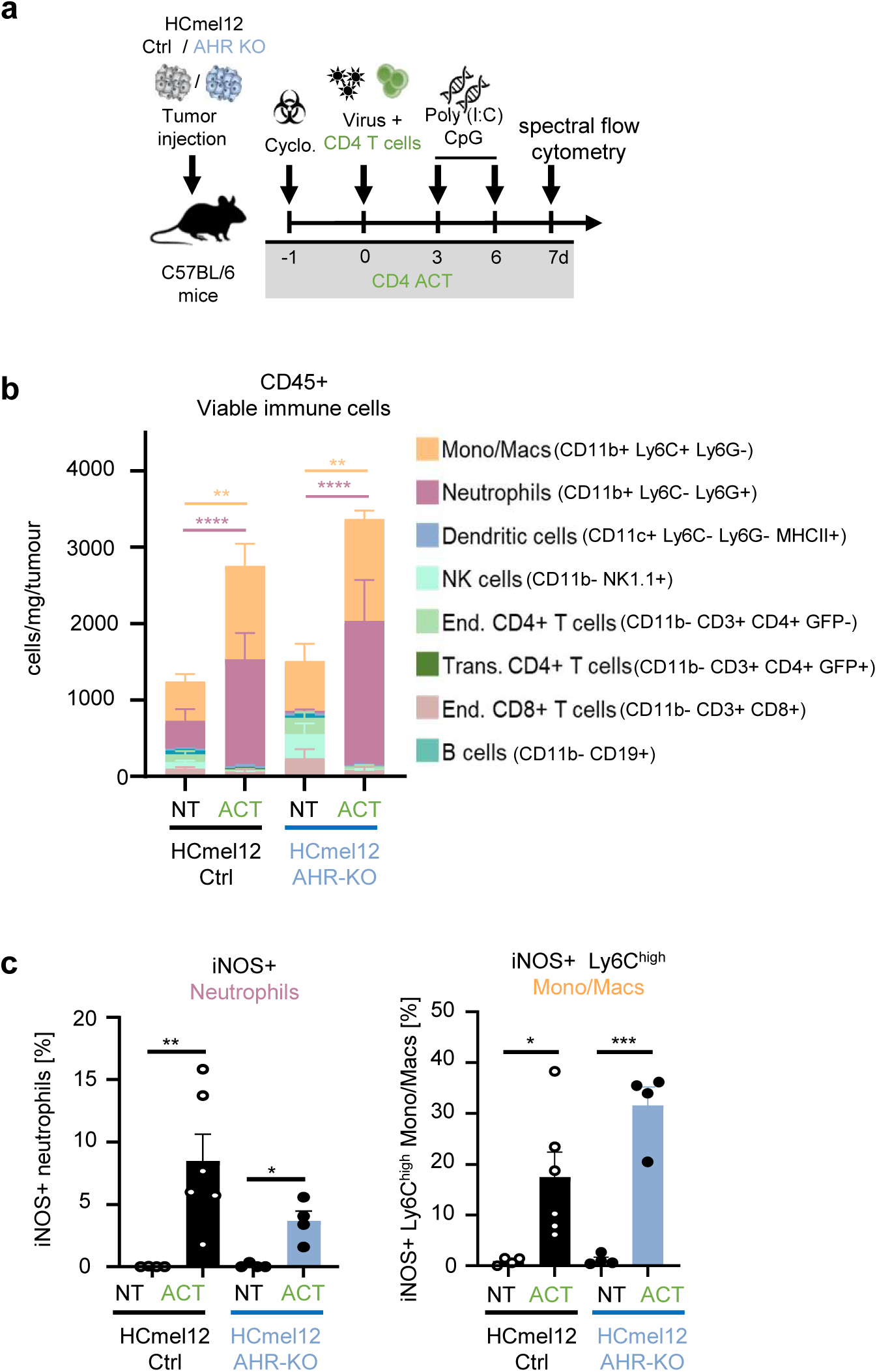
CD4 T-cell immunotherapy promotes infiltration of iNOS-expressing myeloid cells into both AHR-competent and AHR-deficient melanomas. **a**, Experimental protocol for CD4 adoptive T-cell therapy (CD4 ACT) of established HCmel12 CRISPR-control and AHR-KO melanomas in C57BL/6 mice. Tumors were harvested 7 days after CD4+ T-cell transfer for analysis of tumor-infiltrating immune cells by spectral flow cytometry. **b**, Composition and absolute numbers of tumor-infiltrating immune cells in untreated (NT) and CD4 ACT-treated HCmel12 CRISPR-control and AHR-KO melanomas. Stacked bars show the numbers of viable CD45-positive immune cells per milligram of tumor, subdivided into the indicated immune cell populations. **c**, Frequencies of iNOS-expressing neutrophils (left) and Ly6C^high^ monocytes/macrophages (right) among the respective tumor-infiltrating myeloid populations in untreated and CD4 ACT-treated HCmel12 CRISPR-control and AHR-KO melanomas. Bars represent mean ± SD, with individual points indicating biological replicates. Statistical significance was determined using Two-Way ANOVA followed by Tukey’s multiple comparisons test. ns, not significant; \**P* ≤ 0.05; \*\**P* ≤ 0.01; \*\*\**P* ≤ 0.001; \*\*\*\**P* ≤ 0.0001.

## Discussion

Our findings reveal an unexpected role of tumor cell-intrinsic AHR in the response of melanoma to T-cell immunotherapy. Based on our previous observations that AHR promotes inflammation-induced melanoma dedifferentiation and metastatic dissemination (8), we had anticipated that AHR deficiency would reduce resistance to CD8 T-cell therapy targeting melanocytic differentiation antigens (3). Contrary to this expectation, AHR-deficient melanomas exhibited more frequent delayed resistance to CD8 ACT. Our subsequent experiments revealed that AHR-deficient melanoma cells progressively lose IFNγ responsiveness during prolonged inflammatory stimulation, suggesting that AHR may have opposing effects on melanoma immune escape by promoting dedifferentiation while supporting sustained responsiveness to inflammatory signals.

AHR has previously been implicated in the regulation of IFN signaling, including direct interactions with STAT1 (10). In lung adenocarcinoma cells, AHR deficiency was recently shown to impair IFNγ-induced transcription and reduce expression of JAK/STAT signaling components (11). Subsequent work identified additional mechanisms involving the long noncoding RNA INCR1 and the RNA-binding protein HNRNPH1, through which AHR regulates interferon-dependent expression of immune checkpoints and antigen-presentation genes (14). However, the effects of AHR on IFN responses appear to depend on cellular and inflammatory context. In our experiments, AHR-deficient melanoma cells initially exhibited enhanced IFNγ-responsive transcription, consistent with a restraining function of AHR during early stimulation. Surprisingly, prolonged IFNγ exposure resulted in a progressive decline in IFNγ-responsive gene expression and MHC-I and MHC-II surface expression, whereas AHR-competent cells maintained sustained responses. AHR has also recently been identified as a positive regulator of HLA-II expression in human melanoma through direct AHR–ARNT-dependent activation of CIITA independently of IFNγ signaling (15). Thus, AHR may influence antigen presentation through both regulation of IFN responsiveness and direct transcriptional control of antigen-presentation machinery. These findings suggest that tumor cell-intrinsic AHR influences not only the magnitude but also the durability of inflammatory responses, potentially exerting distinct effects during their initiation and maintenance.

The progressive loss of IFNγ responsiveness in AHR-deficient melanoma cells was associated with reduced expression of several intracellular JAK/STAT signaling components, particularly JAK1, whereas expression of the IFNγ receptor genes remained largely preserved. These changes may contribute to the impaired maintenance of IFNγ-responsive transcription, although their causal role remains to be established. Interestingly, treatment with the DNA methyltransferase inhibitor decitabine partially restored MHC expression, raising the possibility that epigenetic mechanisms contribute to the acquired phenotype. Previous studies have demonstrated that prolonged IFN exposure can induce persistent transcriptional and epigenetic changes in tumor cells, including resistance-associated inflammatory programs and dedifferentiation (12,16). In contrast, our findings suggest that AHR deficiency in melanoma cells can lead to a progressive loss of IFNγ-responsive transcription during sustained inflammatory stimulation. Whether the initially enhanced IFNγ response in AHR-deficient melanoma cells triggers counter-regulatory epigenetic changes that subsequently restrict inflammatory responsiveness remains an intriguing question for future investigation.

The acquired loss of IFNγ responsiveness may provide a potential explanation for the increased frequency of delayed resistance to CD8 ACT observed in AHR-deficient melanomas. Tumor cell-intrinsic IFNγ signaling contributes to tumor recognition and elimination by promoting antigen processing and presentation, as supported by experimental studies of T-cell immunotherapy (17), analyses of clinical responses to immune checkpoint blockade in melanoma (18), and observations of acquired resistance to PD-1 blockade in patients with defects in the IFNγ signaling pathway (19). In contrast to inflammation-induced dedifferentiation, which reduces the expression of melanocytic differentiation antigens (3), impaired IFNγ responsiveness may compromise the ability of melanoma cells to maintain antigen-presentation programs during sustained immune pressure. These mechanisms could represent distinct routes of immune escape, with AHR influencing both in opposing directions. However, whether the loss of IFNγ responsiveness observed in vitro directly contributes to delayed CD8 ACT resistance in vivo remains to be established.

Despite their impaired responsiveness to prolonged IFNγ exposure and increased frequency of delayed resistance to CD8 ACT, AHR-deficient melanomas remained susceptible to CD4 T-cell immunotherapy. This finding is consistent with our previous demonstration that CD4 ACT can eradicate melanomas with defective IFNγ signaling through an inflammatory cell death mechanism involving nitric oxide-producing myeloid cells (13). In the present study, AHR-deficient melanoma cells retained susceptibility to the combined cytotoxic effects of a nitric oxide donor and TNFα in vitro, and CD4 ACT induced infiltration of iNOS-expressing myeloid populations into the tumors. These observations suggest that CD4 T-cell-directed inflammatory responses can overcome multiple mechanisms of tumor escape associated with impaired IFNγ responsiveness, although the contribution of nitric oxide to melanoma eradication in the present model remains to be established.

Together, our findings highlight the multifaceted and context-dependent functions of tumor cell-intrinsic AHR in melanoma. While AHR has been implicated in inflammation-induced dedifferentiation and metastatic dissemination, our results reveal an additional role in supporting the maintenance of IFNγ responsiveness during prolonged inflammatory exposure. Although further studies using additional melanoma models and human melanoma cells will be required to establish the broader relevance of these observations, our findings provide important insights into the potential implications of targeting AHR therapeutically, an approach currently being explored in clinical trials (20). In this context, AHR inhibition may offer antitumor benefits while also influencing the capacity of melanoma cells to sustain IFNγ-responsive programs during prolonged immune pressure. Understanding how pharmacological AHR modulation affects these adaptive responses may therefore help refine therapeutic strategies and inform the development of approaches that maximize the benefits of AHR-targeted therapies in combination with immunotherapy.

## Methods

### Mice

Wild-type C57BL/6 mice were obtained from Charles River Laboratories. T-cell receptor-transgenic Pmel-1 (B6.Cg-Thy1ᵃ/Cy Tg(TcraTcrb)8Rest/J) and TRP-1 (B6.Cg-Rag1tm1Mom Tyrp1B-w Tg(Tcra,Tcrb)9Rest/J) mice were originally obtained from The Jackson Laboratory and bred in-house. Pmel-1 mice expressed the congenic marker Thy1.1 (CD90.1). TRP-1 TCR-transgenic mice were crossed onto a congenic CD45.1 (Ly5.1) background. TCR-transgenic T-cell populations were routinely verified by flow cytometry using antibodies against CD90.1 and TCR Vβ13 for Pmel-1 mice and CD45.1 and TCR Vβ14 for TRP-1 mice. The TRP-1 TCR transgene is located on the Y chromosome and is therefore expressed only in male mice. Consequently, all melanoma transplantation and adoptive T-cell therapy experiments were performed using male recipients to avoid potential immune responses against male-specific minor histocompatibility antigens. For the analysis of tumor-infiltrating immune cells, GFP-expressing TRP-1 TCR-transgenic mice, generated by crossing TRP-1 mice with ubiquitously GFP-expressing mice as described previously (13), were used to distinguish transferred from endogenous CD4 T cells.

All mice were maintained under specific pathogen-free conditions in individually ventilated cages at the central animal facility of the University Hospital Magdeburg, Germany, under controlled temperature and humidity conditions and a 12-hour light/dark cycle. Wild-type mice were acclimatized for at least one week before experimental procedures. Unless otherwise indicated, mice were 8–12 weeks old at the beginning of experiments.

All animal experiments were conducted in accordance with Directive 2010/63/EU and German animal welfare legislation (TierSchG and TierSchVersV) and were approved by the competent authority of the State of Saxony-Anhalt, Germany (permit numbers 42502-2-1679 Uni MD, 42502-2-1615 Uni MD, and 42502-2-1672 Uni MD).

### Cell lines and cell culture

The mouse HCmel12 melanoma cell line was established from a primary melanoma in the Hgf-Cdk4^R24C^ mouse model by serial transplantation in our laboratory as described previously (21). AHR-deficient HCmel12 cells were generated by CRISPR/Cas9-mediated gene disruption, together with corresponding AHR-competent CRISPR control cells, as described previously (8). Cells were cultured in complete RPMI 1640 medium (Life Technologies) supplemented with 10% heat-inactivated fetal calf serum (FCS; Biochrom), 1× MEM non-essential amino acids, 1 mM HEPES, 1 mM sodium pyruvate, 2-mercaptoethanol, and penicillin–streptomycin in a humidified incubator containing 5% CO₂. Cell cultures were routinely screened for mycoplasma contamination.

### Tumor transplantation experiments

HCmel12 melanoma cells were harvested during exponential growth at approximately 70–80% confluence, washed, and resuspended in DPBS. A total of 2 × 10⁵ cells in 50 µl DPBS were injected intracutaneously into the shaved right flank of male C57BL/6 mice. Tumor development was monitored by inspection and palpation, and tumor size was measured three times weekly and expressed as mean diameter. Mice were euthanized when tumors exceeded 15 mm mean diameter or when signs of illness necessitated euthanasia according to the approved animal protocols.

### Adoptive cell transfer immunotherapy

Adoptive cell therapy (ACT) was performed as described previously (13). Briefly, when transplanted melanomas reached a mean diameter of 3–5 mm, mice were preconditioned with a single intraperitoneal injection of 2 mg cyclophosphamide (Endoxan, Baxter) in 100 µl PBS one day before T-cell transfer. Splenocytes were isolated from TCR-transgenic Pmel-1 or TRP-1 donor mice containing naïve gp100-specific CD8⁺ or TRP-1-specific CD4⁺ T cells, respectively. Splenocytes containing 5 × 10⁵ Pmel-1 CD8⁺ T cells or 1 × 10⁵ TRP-1 CD4⁺ T cells were transferred intravenously in 100 µl PBS. Transferred T cells were stimulated in vivo by a single intraperitoneal injection of 2.5 × 10^7^ plaque-forming units (PFU) of the recombinant adenoviral vaccine Ad-PT in 100 µl PBS. Ad-PT expresses a fusion protein consisting of the first 150 base pairs of the human *PMEL* cDNA (coding for aa1-50 of the human PMEL/gp100 protein including the CD8+ T-cell epitope KVPRNQDWL recognized by Pmel1 TCR-transgenic T cells) and 1404 base pairs of the mouse *Trp1* cDNA (coding for aa51-518 including the CD4+ T-cell epitope SGHNCGTCRPGWRGAACNQKILTVR recognized by TRP1 TCR-transgenic T cells) followed by sequences coding for a T2A viral self-cleaving peptide and the fluorescent marker protein eYFP (13). On days 3, 6, and 9 after T-cell transfer, tumors were injected with 50 µg CpG oligodeoxynucleotide 1826 (MWG Biotech) and 50 µg polyinosinic:polycytidylic acid (poly(I:C), InvivoGen) in 100 µl distilled water. Seven days after T-cell transfer, peripheral blood was collected from the facial vein to assess expansion of transferred T cells by flow cytometry.

### In vitro chronic IFNγ exposure

HCmel12 AHR-competent and AHR-deficient melanoma cells were cultured in the continuous presence of 1,000 U/ml recombinant mouse IFNγ (PeproTech) for up to 28 days. Twice weekly, cells were harvested, counted, and reseeded in fresh complete medium supplemented with IFNγ at the same concentration. Samples were collected repeatedly from the same cultures at the indicated time points for flow cytometry, RNA sequencing, and other molecular analyses.

### In vitro decitabine treatment

Following 28 days of chronic IFNγ exposure, AHR-competent and AHR-deficient HCmel12 melanoma cells were treated daily with 50 nM decitabine (5-aza-2′-deoxycytidine) or vehicle control and/or IFNγ for 4 days. Melanoma cells that had not undergone the 28-day IFNγ pre-treatment served as controls. Cells were subsequently harvested and analyzed for MHC class I and II surface expression by flow cytometry using an Attune NxT acoustic focusing flow cytometer (Thermo Fisher Scientific).

### In vitro cell death assay

HCmel12 melanoma cells were seeded in flat-bottom 96-well plates in complete RPMI medium. After 4 h, cells were treated with recombinant mouse TNFα (1,000 U/ml, PeproTech) in combination with either recombinant mouse IFNγ (100 U/ml, PeproTech) or the nitric oxide donor S-nitroso-N-acetylpenicillamine (SNAP; 100 µM, Cayman Chemical). After 24 h, floating and adherent cells were collected and stained using the FITC Annexin V Apoptosis Detection Kit I (BD Pharmingen) according to the manufacturer’s instructions. Cell death was assessed by flow cytometry using an Attune NxT acoustic focusing flow cytometer (Thermo Fisher Scientific).

### Flow cytometry

Cultured HCmel12 melanoma cells were harvested and stained with fluorochrome-conjugated monoclonal antibodies according to standard protocols. For analyses of tumor-infiltrating immune cells, excised melanomas were weighed and single-cell suspensions generated by mechanical dissociation with syringe plungers through 70 µm cell strainers. Single suspensions were incubated with anti-CD16/CD32 (1:300, Biolegend) before staining with antibodies against-cell-surface antigens. Intracellular staining was carried out using a Fixation/Permeabilization Solution Kit (BD or Biolegend). Dead cells were excluded using 7-aminoactinomycin D (7-AAD) or fixable viability dyes (Zombie, BioLegend). Flow cytometric measurements were performed using an Attune NxT acoustic focusing flow cytometer (Thermo Fisher Scientific) or a Northern Lights spectral flow cytometer (Cytek Biosciences). Absolute numbers of individual immune cell populations were calculated from their proportions among viable CD45^+^ cells. Antibodies and staining conditions are listed in Supplementary Table 1. Data were analyzed using FlowJo version 10.8.1 for Windows (Waters Biosciences).

### Bulk RNA sequencing

For bulk RNA sequencing, 2 × 10⁵ HCmel12 melanoma cells were collected at the indicated time points during chronic IFNγ exposure. Cells were washed with PBS, pelleted by centrifugation, and stored at −80 °C until processing. RNA isolation, library preparation, and sequencing were performed by Singleron Biotechnologies, targeting a sequencing depth of 20 million reads per sample.

### Bioinformatics and transcriptomic analysis

Read statistics and quality were assessed using SeqKit and FastQC. Sequencing data were processed using the nf-core/rnaseq workflow (v3.14.0) implemented in Nextflow (22), with UMI extraction enabled. Adapters and low-quality bases were trimmed using Trim Galore. Transcript abundances were quantified using Salmon (23) with the *Mus musculus* GRCm39 reference transcriptome (Ensembl release 115) and summarized to gene-level counts using tximport (24).

Gene-level count estimates were used to calculate counts per million (CPM) for expression plots. For differential expression analysis, counts were rounded to integers and restricted to protein-coding genes based on Ensembl gene biotype annotations. Normalization and differential expression analysis were performed using DESeq2 (v1.5.2; 25). Size factors were estimated using the median-of-ratios method to account for differences in library size. Normalized counts were log₂-transformed (log₂[normalized count + 1]) for visualization. Differential expression was assessed using Wald tests for the indicated comparisons, with P- values adjusted using the Benjamini–Hochberg method. Genes with an adjusted P < 0.05 and an absolute apeglm-shrunken log₂ fold change ≥ 1 were considered differentially expressed (26).

Unsupervised k-means clustering was performed in Python using scikit-learn (27). The union of differentially expressed genes across the selected comparisons was filtered to retain genes with a mean normalized count ≥ 50 in at least one condition. For each gene, log₂-transformed normalized counts were averaged across biological replicates within each condition and standardized to Z-scores across conditions. K-means clustering was performed with random_state = 0 and n_init = 10. The number of clusters was selected using the elbow method (within-cluster sum of squares for k = 1–15) and visual inspection. Cluster centroids were calculated as the mean Z-score profiles of the corresponding genes.

Each cluster was analyzed for overrepresentation of the 50 MSigDB Hallmark gene sets (28), using the native *Mus musculus* gene sets (release 2026.1; 29). All genes tested in the differential expression analysis served as the background, and gene sets containing 5–2,000 background genes were included. Enrichment was assessed using a one-sided hypergeometric test implemented in SciPy. P-values were adjusted using the Benjamini– Hochberg method, and gene sets with a false discovery rate (FDR) < 0.05 were considered significantly enriched.

### Western blot analysis

HCmel12 melanoma cells were lysed using M-PER Mammalian Protein Extraction Reagent supplemented with protease inhibitors (Thermo Fisher Scientific). Protein concentrations were determined using the Pierce BCA Protein Assay Kit (Thermo Fisher Scientific). Lysates containing 10 µg protein were mixed with Laemmli sample buffer, heated at 95 °C for 5 min, and separated by SDS-PAGE using 10% polyacrylamide gels. Proteins were transferred onto 0.2 µm PVDF membranes by wet blotting. Membranes were blocked with 5% skimmed milk in Tris-buffered saline containing Tween 20 (TBST) for 1 h and incubated overnight at 4 °C with a primary rabbit anti-mouse JAK1 antibody (Cell Signalling clone 6G4) or anti mouse β-actin antibody (Santa Cruz clone C4) as a loading control. Bound antibodies were detected using the appropriate horseradish peroxidase-conjugated secondary antibody (Cell Signalling) followed by SignalFire ECL reagent (Cell Signaling Technology) and visualization with an Octoplus QPLEX imager (NH DyeAgnostics).

### Statistical analysis

Survival data were analyzed using the Kaplan–Meier method. Differences between survival curves were assessed using the log-rank (Mantel–Cox) test. For other experimental data, statistical differences between groups were assessed using either one-way or two-way analysis of variance (ANOVA), as appropriate for the experimental design. One-way ANOVA was used when comparing a single independent variable across multiple groups, whereas two-way ANOVA was applied when two independent variables were investigated simultaneously, allowing for the assessment of both individual factor effects and potential interactions between them. Where significant differences were detected, Tukey’s multiple comparisons test was performed for post hoc pairwise comparisons between groups while controlling for multiple testing. All statistical tests were two-sided, and differences were considered statistically significant at p < 0.05.

## Data availability

The RNA-seq data generated in this study will be deposited in the NCBI Gene Expression Omnibus (GEO) repository and made publicly available upon publication of the peer-reviewed manuscript. Additional data supporting the findings of this study are available from the corresponding author upon reasonable request.

## Acknowledgements

We would like to thank K. Beinhoff, J. Herz, and J. Leipold for their assistance with mouse colony management, mycoplasma testing of cell cultures, and genotyping and phenotyping of the mouse colonies. We are also grateful to Martin Böttcher and Dimitrios Mougiakakos for providing access to the spectral flow cytometer and for their assistance with data acquisition. T.T. was supported by funding from the German Research Foundation (FOR5489-P8) and the German Cancer Aid (grant nos. 70112525 and 70114549). T.H.S. was supported by funding from the German Research Foundation (FOR5489-P7).

## Author contributions

S.G., W.L., K.K., S.B., S.W. performed experiments and analysed data. M.M. generated cell lines. S.B. and B.K. generated recombinant adenoviruses. A.D.B. and M.C. performed bioinformatic analyses. S.G., B.K., A.C.B., and T.T. designed experiments. S.G., B.K., A.C.B., A.D.B., E.G., T.H.S. and T.T. contributed intellectual input and helped to interpret data. T.T. led the research program. S.G., M.C., T.H.S., E.G. and T.T. wrote the manuscript.

## References

1. Wolchok JD, Chiarion-Sileni V, Rutkowski P, et al. Final, 10-year outcomes with nivolumab plus ipilimumab in advanced melanoma. N Engl J Med. 2025;392:11–22. 10.1056/NEJMoa2407417

2. Robertson BM, Fane ME, Weeraratna AT, et al. Determinants of resistance and response to melanoma therapy. Nat Cancer. 2024;5:964–982. 10.1038/s43018-024-00794-1

3. Landsberg J, Kohlmeyer J, Renn M, et al. Melanomas resist T-cell therapy through inflammation-induced reversible dedifferentiation. Nature. 2012;490:412–416. 10.1038/nature11538

4. Opitz CA, Holfelder P, Prentzell MT, Trump S. The complex biology of aryl hydrocarbon receptor activation in cancer and beyond. Biochem Pharmacol. 2023;216:115798. 10.1016/j.bcp.2023.115798

5. Coumoul X, Barouki R, Esser C, et al. The aryl hydrocarbon receptor: structure, signaling, physiology and pathology. Signal Transduct Target Ther. 2026;11:20. 10.1038/s41392-025-02500-8

6. Corre S, Tardif N, Mouchet N, et al. Sustained activation of the aryl hydrocarbon receptor transcription factor promotes resistance to BRAF-inhibitors in melanoma. Nat Commun. 2018;9:4775. 10.1038/s41467-018-06951-2

7. Contador-Troca M, Alvarez-Barrientos A, Barrasa E, et al. The dioxin receptor has tumor suppressor activity in melanoma growth and metastasis. Carcinogenesis. 2013;34:2683– 2693. 10.1093/carcin/bgt248

8. Mengoni M, Braun AD, Gaffal E, Tüting T. The aryl hydrocarbon receptor promotes inflammation-induced dedifferentiation and systemic metastatic spread of melanoma cells. Int J Cancer. 2020;147:2902–2913. 10.1002/ijc.33252

9. Kohlmeyer J, Cron M, Landsberg J, et al. Complete regression of advanced primary and metastatic mouse melanomas following combination chemoimmunotherapy. Cancer Res. 2009;69:6265–6274. 10.1158/0008-5472.CAN-09-0579

10. Kimura A, Naka T, Nohara K, Fujii-Kuriyama Y, Kishimoto T. Aryl hydrocarbon receptor regulates Stat1 activation and participates in the development of Th17 cells. Proc Natl Acad Sci USA. 2008;105:9721–9726. 10.1073/pnas.0804231105

11. Snyder M, Wang Z, Lara B, et al. The aryl hydrocarbon receptor controls IFN-γ-induced immune checkpoints PD-L1 and IDO via the JAK/STAT pathway in lung adenocarcinoma. J Immunol. 2025;214:413–432. 10.1093/jimmun/vkae023

12. Benci JL, Xu B, Qiu Y, et al. Tumor interferon signaling regulates a multigenic resistance program to immune checkpoint blockade. Cell. 2016;167:1540–1554.e12. 10.1016/j.cell.2016.11.022

13. Kruse B, Buzzai AC, Shridhar N, et al. CD4+ T cell-induced inflammatory cell death controls immune-evasive tumours. Nature. 2023;618:1033–1040. 10.1038/s41586-023-06199-x

14. Lara B, Snyder M, Fimbres J, et al. The AhR regulates IFN-induced immune checkpoints in lung cancer cells through HNRNPH1, an RNA-binding protein, and INCR1, a novel long non-coding RNA. J Biol Chem. 2025;301:110316. 10.1016/j.jbc.2025.110316

15. Jin Y, Zheng W, Zhang R, et al. The aryl hydrocarbon receptor (AHR) drives human leukocyte antigen (HLA)-II expression in human melanoma. J Exp Clin Cancer Res. 2026;45:78. 10.1186/s13046-026-03673-y

16. Kim YJ, Sheu KM, Tsoi J, et al. Melanoma dedifferentiation induced by IFN-γ epigenetic remodeling in response to anti-PD-1 therapy. J Clin Invest. 2021;131:e145859. 10.1172/JCI145859

17. Patel SJ, Sanjana NE, Kishton RJ, et al. Identification of essential genes for cancer immunotherapy. Nature. 2017;548:537–542. 10.1038/nature23477

18. Grasso CS, Tsoi J, Onyshchenko M, et al. Conserved interferon-γ signaling drives clinical response to immune checkpoint blockade therapy in melanoma. Cancer Cell. 2020;38:500– 515.e3. 10.1016/j.ccell.2020.08.005

19. Zaretsky JM, Garcia-Diaz A, Shin DS, et al. Mutations associated with acquired resistance to PD-1 blockade in melanoma. N Engl J Med. 2016;375:819–829. 10.1056/NEJMoa1604958

20. Aggen DH, McKean M, Hoffman-Censits JH, et al. IK-175, an oral aryl hydrocarbon receptor inhibitor, alone and with nivolumab in patients with advanced solid tumors and urothelial carcinoma. Clin Cancer Res. 2026;32:94–105. 10.1158/1078-0432.CCR-25-2013

21. Bald T, Quast T, Landsberg J, et al. Ultraviolet-radiation-induced inflammation promotes angiotropism and metastasis in melanoma. Nature. 2014;507:109–113. 10.1038/nature13111

22. Ewels PA, Peltzer A, Fillinger S, et al. The nf-core framework for community-curated bioinformatics pipelines. Nat Biotechnol. 2020;38:276–278. 10.1038/s41587-020-0439-x

23. Patro R, Duggal G, Love MI, Irizarry RA, Kingsford C. Salmon provides fast and bias-aware quantification of transcript expression. Nat Methods. 2017;14:417–419. 10.1038/nmeth.4197

24. Soneson C, Love MI, Robinson MD. Differential analyses for RNA-seq: transcript-level estimates improve gene-level inferences. F1000Research. 2015;4:1521. 10.12688/f1000research.7563.2

25. Love MI, Huber W, Anders S. Moderated estimation of fold change and dispersion for RNA-seq data with DESeq2. Genome Biol. 2014;15:550. 10.1186/s13059-014-0550-8

26. Zhu A, Ibrahim JG, Love MI. Heavy-tailed prior distributions for sequence count data: removing the noise and preserving large differences. Bioinformatics. 2019;35:2084–2092. 10.1093/bioinformatics/bty895

27. Pedregosa F, Varoquaux G, Gramfort A, et al. Scikit-learn: machine learning in Python. J Mach Learn Res. 2011;12:2825–2830. https://dl.acm.org/doi/10.5555/1953048.2078195

28. Liberzon A, Birger C, Thorvaldsdóttir H, et al. The Molecular Signatures Database (MSigDB) hallmark gene set collection. Cell Syst. 2015;1:417–425. 10.1016/j.cels.2015.12.004

29. Castanza AS, Recla JM, Eby D, et al. Extending support for mouse data in the Molecular Signatures Database (MSigDB). Nat Methods. 2023;20:1619–1620. 10.1038/s41592-023-02014-7

