## Supplemental Table 1 for "Tumor cell-intrinsic AHR sustains IFNγ responsiveness and durable CD8 T-cell-mediated melanoma control"

**Supplementary Table 1. Antibodies for flow cytometry analyses**

***Antibodies used to analyze the effect of chronic IFNγ on Hcmel12 melanoma cells***

| Antigen | Fluorophore | Clone | Dilution | Manufacturer |
|---|---|---|---|---|
| Anti-mouse CD119 (Ifngr1) | BV421 | GR20 | 1:1600 | BD Bioscience |
| Anti-mouse MHCI (H2kb) | PeCy7 | AF6-88.5 | 1:4000 | Biolegend |
| Anti-mouse MHCII (I-A/I-E) | BV510 | M5/114.15.2 | 1:800 | Biolegend |
| Anti-mouse PD-L1 | PE | 10F.9G2 | 1:800 | Biolegend |
| Live/dead | Annexin V FITC | - | 1:100 | Sigma Aldrich |
| Live/dead | Propidium Iodide | - | 1:100 | Sigma Aldrich |

***Antibodies used to detect TCR-transgenic T cells for ACT experiments***

| Antigen | Fluorophore | Clone | Dilution | Manufacturer |
|---|---|---|---|---|
| Anti-mouse CD3 | BV421 | 17A2 | 1:400 | Biolegend |
| Anti-mouse CD4 | PE | GK1.5 | 1:1600 | ThermoFisher |
| Anti-mouse CD8a | APC | 53-6.7 | 1:600 | Biolegend |
| Anti-mouse CD45.2 | APC | 104 | 1:200 | BD Bioscience |

***Antibodies used for analysis of tumor-infiltrating immune cells***

| **Antigen** | **Fluorophore** | **Clone** | **Dilution** | **Manufacturer** |
|---|---|---|---|---|
| Anti-mouse CD3 | BV785 | 17A2 | 1:50 | Biolegend |
| Anti-mouse CD4 | BV605 | RM-4-5 | 1:500 | BD Bioscience |
| Anti-mouse CD8 | VioGreen | 53-6.7 | 1:100 | Miltenyi |
| Anti-mouse CD11b | BV711 | M1/70 | 1:200 | Biolegend |
| Anti-mouse CD11c | APC-Fire750 | N418 | 1:200 | Biolegend |
| Anti-mouse CD16/32 | none | 93 | 1:300 | Biolegend |
| Anti-mouse CD19 | PE-fire640 | 6D5 | 1:400 | Biolegend |
| Anti-mouse CD44 | BV421 | IM7 | 1:1600 | BD Bioscience |
| Anti-mouse CD45 | PerCP | 30-F11 | 1:250 | BD Bioscience |
| Anti-mouse CD80 | BV750 | 16-10A1 | 1:400 | BD Bioscience |
| Anti-mouse CD206 | BV650 | C068C2 | 1:200 | Biolegend |
| Anti-mouse CX3CR1 | APC-Fire810 | SA011F11 | 1:100 | Biolegend |
| Anti-mouse CXCR3 | BV480 | CXCR3-173 | 1:400 | BD Bioscience |
| Anti-mouse F4/80 | APC | BM8 | 1:200 | Biolegend |
| Anti-mouse iNOS | PE | CXNFT | 1:300 | ThermoFisher |
| Anti-mouse Ly6C | R718 | AL-21 | 1:200 | BD Bioscience |
| Anti-mouse Ly6G | PE-efluor610 | 1A8-Ly6g | 1:1600 | ThermoFisher |
| Anti-mouse MHCII (I-A/I-E) | PerCP-eFluor710 | M5/114.15.2 | 1:1600 | ThermoFisher |
| Anti-mouse NK1.1 | BV570 | PK136 | 1:200 | Biolegend |
| anti-mouse PD-1 | PE-Fire810 | 29F.1A12 | 1:200 | Biolegend |
| Anti-mouse PD-L1 | AF660 | MIH5 | 1:400 | ThermoFisher |
| Anti-mouse Siglec H | PE-Vio770 | REA819 | 1:400 | Miltenyi |
| Live/dead | Zombie NIR | - | 1:5000 | Biolegend |
